# Continuous attractor circuits for decision making with Laplace-domain neural representations

**DOI:** 10.64898/2026.08.03.742594

**Authors:** Chenyu Wang, Rui Cao, Marc W. Howard

## Abstract

Decision formation is commonly described as the accumulation of noisy evidence in a low-dimensional decision variable, but it remains unclear how this latent computation is implemented by heterogeneous neural responses. Here, we propose that ramping and sequentially firing neurons form complementary population codes for the same decision variable. Inspired by Laplace-domain neural representations of time, exponential receptive fields in a ramping population give rise to a translatable edge-like activity profile; localized receptive fields in a sequential population give rise to an aligned bump-like profile. We construct a continuous attractor neural network that dynamically maintains these complementary representations while implementing evidence accumulation along a shared latent manifold. At the behavioral level, simulations show that the circuit closely reproduces the single-trial trajectories, choice probabilities, and reaction-time statistics of a standard diffusion decision model while generating heterogeneous ramping and sequential neural responses. Our framework connects latent behavioral dynamics, population geometry, and recurrent circuit mechanisms. More broadly, it provides a circuit-level realization of computation in the Laplace domain that may support the representation and updating of continuous cognitive variables across decision making, timing, memory, and spatial cognition.

## Introduction

Evidence accumulation tasks provide a central experimental paradigm for studying how organisms form decisions under uncertainty. In these tasks, subjects must integrate noisy sensory information over time before committing to one of several alternatives, making it possible to relate stimulus statistics, choice accuracy, and reaction times to an underlying latent decision process. Diffusion decision models (DDM) and related sequential-sampling models have therefore become a standard framework for this problem. DDMs capture evidence integration in terms of a low-dimensional decision variable evolving toward decision bounds. These models successfully account for key behavioral signatures such as psychometric curves, response-time distributions, and speed–accuracy tradeoffs [1–3].

Many studies have sought to identify neural markers of such a latent decision variable. Early work established a general framework in which decision formation is reflected in neural activity that integrates evidence during deliberation and approaches commitment over time [4, 5]. Such ramping activity was famously characterized in area LIP of posterior parietal cortex during primate motion-discrimination tasks, with related buildup signals reported in oculomotor circuits such as frontal eye field and superior colliculus [4–6]. More recent rodent studies showed that accumulation-related ramping dynamics are distributed across multiple regions, including posterior parietal cortex (PPC) [7–9], frontal orienting fields (FOF) [7–9], and anterior dorsal striatum (ADS) [9–12], with these responses unfolding over diverse timescales and reflecting region-specific contributions to accumulated evidence and choice formation [8, 9, 11]. However, subsequent population-level studies suggested that this ramping picture captures only part of a richer dynamical organization. In mouse PPC, decision-related neurons can fire transiently and form choice-specific sequences across the trial [13–15]. Related work in primate dorsal premotor cortex (PMd) further showed that, during perceptual decisions, ramping-like and sequence-like single-neuron responses can coexist within the same population and reflect nonlinear tuning to a shared low-dimensional decision variable [16]. Together, these observations point to a broader heterogeneity of decision-related neural dynamics, spanning not only diverse contributions of different brain regions to the same evidence accumulation process, but also distinct temporal motifs such as ramping and sequential activity.

A natural way to interpret this heterogeneity is to view decision-related neural activity as different neural representations of a shared latent decision variable. This perspective has been formalized in models that jointly infer latent decision dynamics and neural activity, while characterizing individual neurons by their tunings to the latent decision variable [9,16]. Yet, it remains unclear how ramping-like and sequential firing patterns can emerge in a principled way as distinct response motifs of the same latent variable, and how such heterogeneity could arise from an underlying circuit mechanism.

Inspired by previous work on Laplace-domain neural representations of time [17], we address this challenge by proposing that evidence accumulation under drift-diffusion dynamics is represented through complementary population codes over a latent decision variable [18]. Ramping receptive fields arise from a Laplace transform of the latent decision variable with different real constants. Sequential receptive fields arise from the corresponding inverse Laplace transform of the same variable. We show that the two motifs—ramping activity and sequential activity—can be interpreted as aligned edge and bump manifolds over the same decision variable. Building on recent continuous-attractor implementations of Laplace neural manifolds [19], we construct a coupled continuous attractor circuit composed of an edge-like manifold for ramping neurons and a bump-like manifold for sequentially firing neurons. By tuning the reciprocal coupling between the two populations, the model implements a shared latent decision variable with diffusion-like dynamics, while flexibly expressing it as ramping or sequential activity in individual neurons. We show that this model reproduces DDM-like choice and reaction-time behavior while predicting heterogeneous neural responses organized into aligned ramping and sequential population codes. Our results reveal how complementary Laplace-domain representations can be coupled in a recurrent circuit model to implement diffusion-like decision dynamics while generating diverse neural response motifs.

## Results

### Theory: Population geometries of ramping and sequential neurons

We consider neural tuning in a two-choice evidence-accumulation task. We assume two choice-selective groups of neurons, each preferentially responsive to one of the two decision bounds. Within each choice-selective group, we consider two functionally distinct populations: one composed of ramping neurons and the other of sequentially firing neurons. Each functional population consists of *N* neurons sharing the bound preference of its choice-selective group and embedded in a one-dimensional neural space, with coordinates *θ*_*i*_ sampled independently and uniformly from a finite interval *C* = [*θ*_min_, *θ*_max_] (Fig. 1C, left). The activity of each neuron is determined by a neuron-specific tuning function of an internally generated one-dimensional decision variable evolving according to a standard DDM with symmetric absorbing boundaries (Fig. 1A). To describe the tuning of both groups within a unified framework, we express this decision variable using a common distance-to-bound convention: for each group, *x*(*t*) ∈ [0, 1] denotes the normalized distance of the latent decision state from that group’s preferred bound, with *x* = 0 at the preferred bound and *x* = 1 at the opposite bound. Under this convention, the same tuning-function framework can be used to characterize both choice-selective groups.

**Figure 1:**
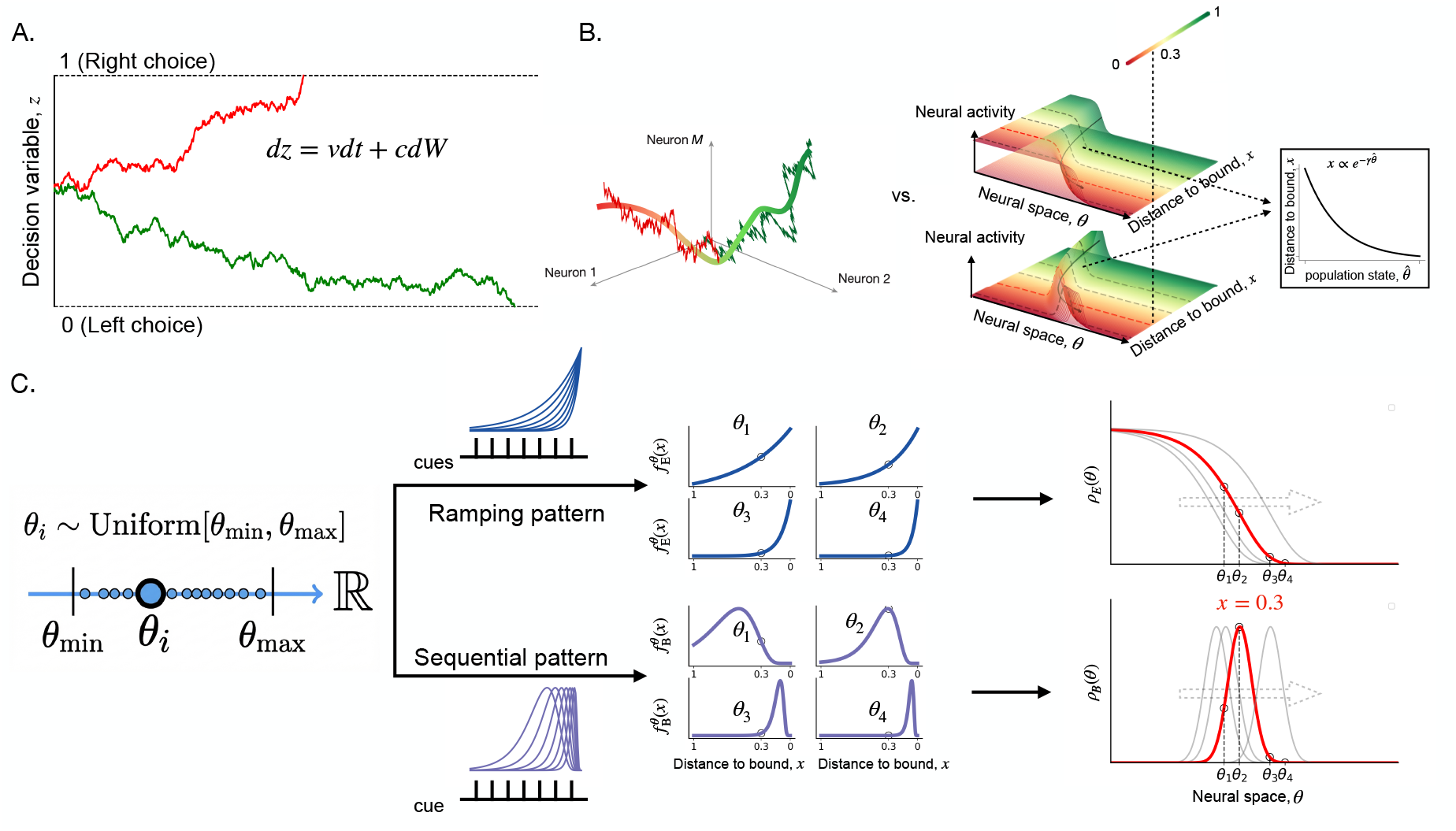
Geometric population representations of a decision variable. **(A)** Schematic of a classical diffusion decision process in which a one-dimensional decision variable *x*(*t*) evolves stochastically between symmetric absorbing boundaries corresponding to left (*x* = 1) and right (*x* = 0) choices. (**B**) Comparison between a general geometric coding principle and the coding principle proposed here. In the general framework (left, from [16]), neural population responses evolve along a one-dimensional manifold that encodes the latent decision variable. Here, we propose a more specific population code in which two aligned populations, edge and bump, form a translationally invariant activity profile, and the decision variable is encoded by the position of edge or bump through a nonlinear mapping. This population geometry arises naturally from the assumed tuning rules for ramping and sequentially firing neurons (fig. 1C). The rightmost inset illustrates the nonlinear mapping between the shared edge–bump position 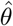 in neural space and the represented decision variable *x*. Consequently, equal displacements along the neural coordinate correspond to position-dependent changes in *x*. **(C)** Construction of ramping (edge) and sequential (bump) receptive fields in neural space. Neurons are embedded along a one-dimensional neural coordinate *θ* (left), and each neuron has a receptive field over the distance-to-bound variable *x* (middle). Ramping neurons are modeled with exponential receptive fields (see (1)) whose rate constants vary systematically with neural position (see (2)), whereas sequential neurons have localized receptive fields (see (8)). In the continuum limit, the organization of these receptive fields across *θ* gives rise to smooth population activity profiles in neural space (right): an edge-like profile *ρ*_*E*_(*θ*) for the ramping population and a localized bump *ρ*_*B*_(*θ*) for the sequential population. Conversely, translation of the edge profile across neural space gives rise to ramping receptive fields over *x*, whereas translation of the bump profile gives rise to sequential receptive fields. The circle markers denote the four example units in the middle panel. The gray edge and bump profiles represent the same family of population states shown in gray in Fig. 1B (right), whereas the red edge-bump state marks the highlighted example at *x* = 0.3.

#### Ramping neurons form an edge population representation

The firing rate of the i-th ramping neuron is assumed to depend on *x* through an exponential tuning function

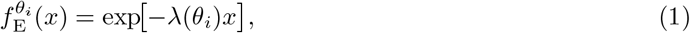

where we define the ramping-rate function *λ*(·) to vary across *θ* following a geometric progression,

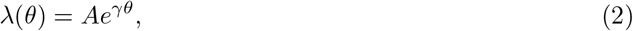

where *A* > 0 sets the overall scale of the ramping rates and *γ* controls how rapidly these rates change across neural space *θ*. The upper middle of Fig. 1C shows examples of these exponential tuning functions with different ramping rates. In the continuum neural field limit (*N* → ∞), and for pre-boundary states *x* ∈ [*x*_min_, *x*_max_] ⊂ [0, 1] the population response of ramping neurons can be described by a continuous function of *θ*,

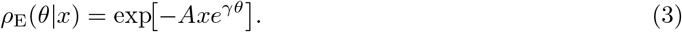

That is, the population activity takes a double-exponential form in *θ*. This double-exponential structure produces an intrinsic edge-like geometry in *θ*, characterized by a rapid transition from saturation to suppression at a location controlled by evidence *x* (Fig. 1C, upper right). The lower cutoff *x*_min_ > 0 excludes the degenerate bound state *ρ*_E_(*θ* | 0) = 1, at which the population no longer has an edge. *A* sets the overall scale and *γ* controls how sharply the edge is organized across neural space *θ*. Crucially, changes in *x* do not deform this geometry; instead, they induce a rigid translation of the entire population profile along *θ*. To make this invariance explicit, we characterize the population profile by the location of its edge, defined as the point at which the response undergoes its most rapid transition from saturation to suppression, 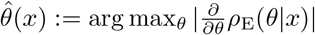. This edge location admits a closed-form expression,

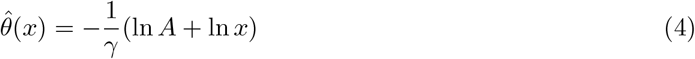

revealing that the decision variable is encoded logarithmically as a phase shift of the edge along the *θ*-manifold. We thereby rewrite *ρ*_E_ as:

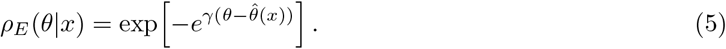

For analytical tractability, we approximate this double-exponential edge profile by a logistic sigmoid, which preserves the sharp transition around 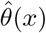 while yielding more convenient analytic expressions (e.g., for derivatives and inverse mappings):

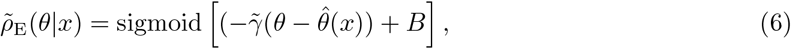

where we choose 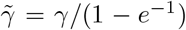 and *B* = − log(*e* − 1) to match the local slope of the target edge profile (see Appendix). We use this sigmoid approximation in the following analysis.

#### Sequentially firing neurons form a bump population representation

Sequentially firing neurons form a localized bump of activity at the population level, analogous to canonical continuous attractor representations such as head-direction cells [20–22] and place cells [23– 25]. Crucially, this sequential pattern is also nonlinear: population activity progresses faster as the decision variable approaches commitment, mirroring the same logarithmic warping implied by the ramping population’s dynamics (see (4)). We therefore model the population response of bump neurons by a Gaussian function whose center follows the same decision-variable–dependent dynamics as the edge (Fig. 1C, lower right),

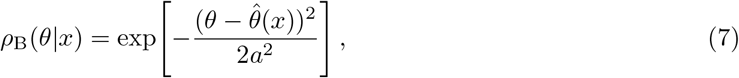

where *a* > 0 controls the spatial width of the bump along the neural coordinate *θ*.

At the individual level, the firing rate of a sequentially firing neuron at *θ*_*i*_ is obtained by evaluating the population bump profile at that fixed neural coordinate. Because the bump center 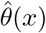 depends linearly on ln *x*, each neuron has a Gaussian tuning curve in log-*x* space. We therefore require the single-neuron tuning curve of bump neurons as

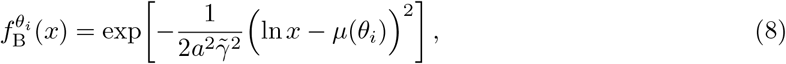

where the location parameter *µ*(*θ*_*i*_) satisfies

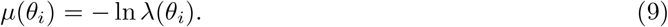

Together, these results show that changes in the decision variable *x* induce rigid translations of both the ramping (edge) and sequentially firing (bump) population responses along neural space, yielding aligned representations of the same latent decision state (Fig. 1C, right). This construction provides a specific realization of the general geometric coding principle illustrated in Fig. 1B. In the general framework, the latent decision variable parameterizes neural population activity along a one-dimensional manifold (Fig. 1B, left; [16]). Here, the single-neuron tuning structure further specifies the geometry of this manifold: an edge and a bump with fixed shapes remain aligned and translate together along the neural coordinate *θ*, with their shared position related nonlinearly to *x* through 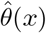 (Fig. 1B, right). Thus, the proposed tuning functions determine not only heterogeneous single-neuron response motifs, but also an explicit population-level geometry for representing the latent decision variable.

### A Biologically plausible mechanism for decision-variable encoding via Continuous attractor neural network

Here we construct a circuit-level model whose dynamics are explicitly designed to implement the population geometries described above (Fig. 2A, B). Our goal is to realize a coupled edge–bump state in which the edge and bump remain aligned and translate together along the neural coordinate *θ*, so that both populations encode the same latent decision state at every moment. Through the mapping 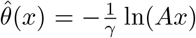, this shared position provides a logarithmically warped coordinate for the decision variable. We refer to this architecture as the edge–bump CANN model.

**Figure 2:**
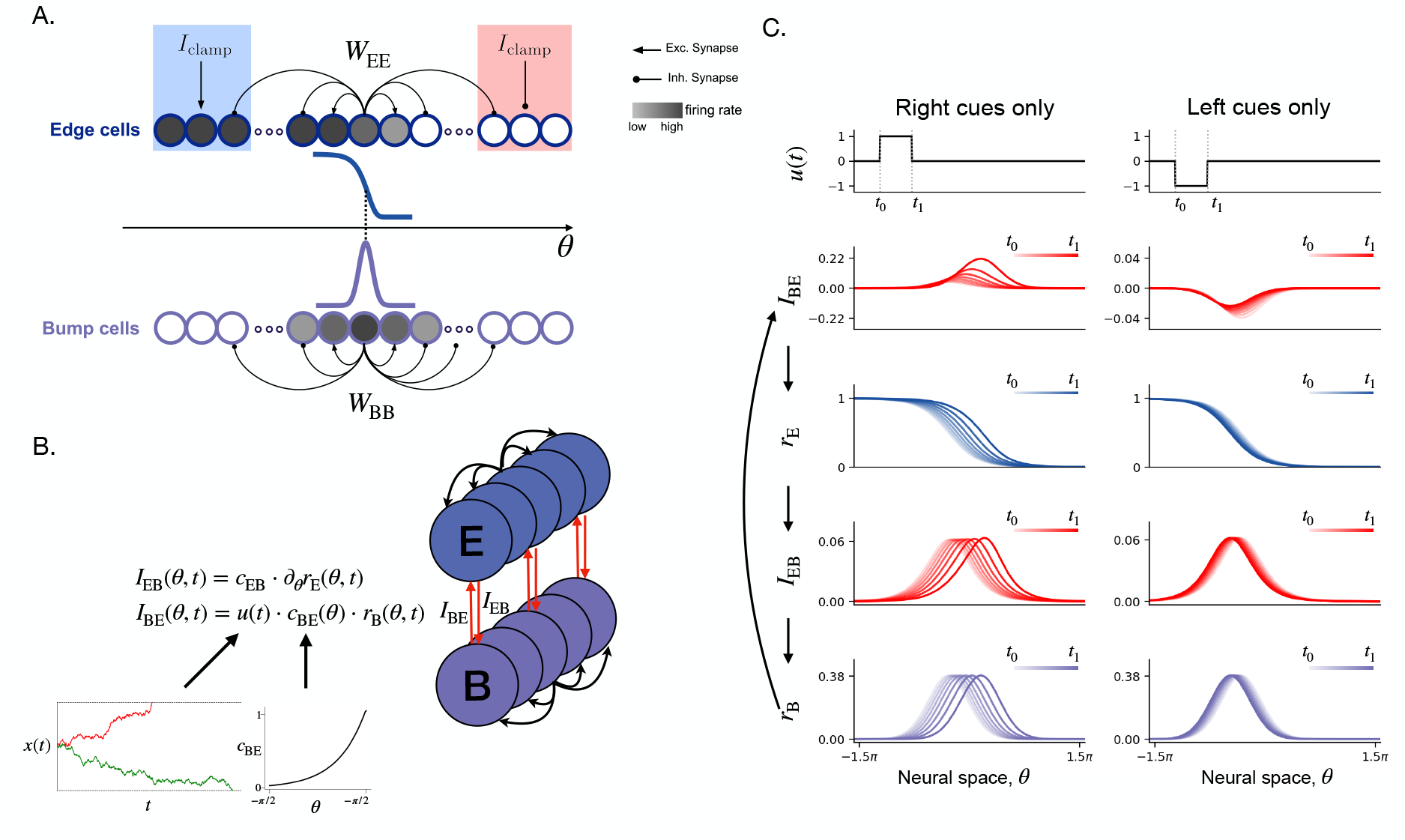
A CANN mechanism for the edge and bump population codes. **(A)** Schematic of the recurrent edge and bump populations that implement the edge and bump geometries introduced in Fig. 1C. In the edge population, recurrent interactions *W*_EE_ together with the boundary-stabilizing input *I*_clamp_ stabilize an edge state whose position *θ*(*t*) encodes the decision variable. In the bump population, recurrent interactions *W*_BB_ stabilize a localized bump state. The circles are shaded from gray to white to indicate firing rate from high to low. **(B)** Circuit architecture of the coupled CANN model. The edge-to-bump pathway aligns the edge and bump states, whereas the bump-to-edge input *I*_BE_ = *u*(*t*)*c*_BE_(*θ*)*r*_B_ drives their translation along the attractor manifold. Here, *u*(*t*) specifies the target latent dynamics, while *c*_BE_(*θ*) corrects for the nonlinear mapping 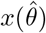 shown in the inset. (**C**) Simulated dynamics of the coupled CANN under deterministic drive. This panel shows direct simulations of the model, with *u*_noise_ = 0 and only the drift term *u*_drift_ retained in the bump-to-edge input. The left column shows a constant positive drive (*u*_drift_ > 0), and the right column shows a constant negative drive (*u*_drift_ < 0). From top to bottom, the rows show the input drive, the resulting bump-to-edge input *I*_BE_, the translated edge state *r*_E_, the induced edge-to-bump input *I*_EB_, and the tracked bump state *r*_B_, plotted over time and neural space. The arrows highlight the closed dynamical loop *I*_BE_ → *r*_E_ → *I*_EB_ → *r*_B_ → *I*_BE_: the bump-to-edge input *I*_BE_ shifts the edge state *r*_E_, the shifted edge generates an edge-to-bump input *I*_EB_, this localized input pulls the bump state *r*_B_ toward alignment with the edge, and the resulting bump state feeds back to the bump-to-edge pathway. To ensure that a constant drift produces linear dynamics in decision-variable space, the mapping from the decision variable to neural space is specified to be nonlinear (Eq. 4). Consequently, in neural space the coupled edge–bump state moves rightward with increasing speed for constant positive drive and leftward with decreasing speed for constant negative drive.

#### Continuous Attractor Circuit for Edge-Bump Geometry

We model our neural circuit by standard rate-based neurons with recurrent network connectivity and point-wise neural nonlinearity. Consider two interacting neural populations: an edge population, corresponding to ramping neurons, and a bump population, corresponding to sequentially firing neurons. In each population, neurons are embedded in a one-dimensional Euclidean coordinate space, with co-ordinates *θ*_*i*_ sampled uniformly from a finite interval. The firing rate of a neuron at coordinate *θ*_*i*_ is denoted by *r*_E_(*θ*_*i*_) or *r*_B_(*θ*_*i*_), respectively. For analytical tractability, and to allow synaptic weights and neural activations to be expressed as functions instead of finite-dimensional vectors, we adopt the continuum neural field limit. The recurrent edge and bump architectures are illustrated schematically in Fig. 2A. The construction below first establishes aligned edge and bump attractors parameterized by this shared coordinate, and then specifies reciprocal coupling so that incoming evidence translates the aligned state by the appropriate amount in latent decision space. The network dynamics are given by:

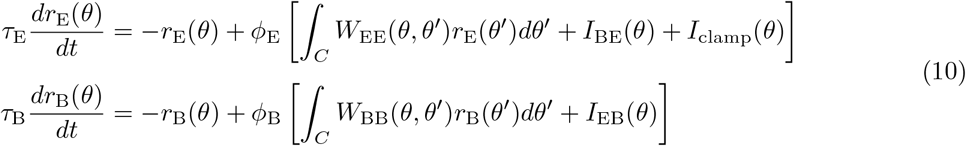

where *τ*_E_ and *τ*_B_ are the time constants of the edge and bump populations, *W*_EE_ and *W*_BB_ denote recurrent connectivity within each population, *I*_BE_ and *I*_EB_ denote the reciprocal bump–edge input currents, and *ϕ*_E_, *ϕ*_*B*_ are the nonlinearities of each population. The extra term *I*_clamp_(*θ*) is a boundary-stabilizing input current that anchors the edge state on the finite interval (see Fig. 2A). For the complete model specification, see Methods.

Our first goal is to identify interaction weights *W*_BB_ and *W*_EE_ that support stable bump and edge attractor states given by (6) and (7). This family of attractor states provides a continuous neural representation of the decision variable *x*, such that changes in *x* correspond to translations of the bump/edge profile along the *θ*-manifold. To remain consistent with previous work on CANNs [20, 26, 27], we assume *W*_BB_ and *W*_EE_ to be functions of distance between neurons in coordinate space:

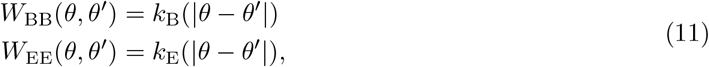

where *k*_B_ and *k*_E_ are even functions of neural distance, preserving translational symmetry in the interior of the neural domain. The two kernels are chosen for different reasons. For the bump population, we use a Gaussian kernel as it admits an analytically solvable family of Gaussian stationary states that exactly matches the desired bump profile *ρ*_B_. More general center–surround kernels could also support bump-like states, but would not provide the same exact Gaussian solution. For the edge population, no analogous closed-form construction is available for the desired sigmoidal profile *ρ*_E_. We therefore use a difference-of-Gaussians kernel as a flexible, translation-invariant family with independently tunable local excitation and broader inhibition. Its parameters are calibrated so that the target edge profile is approximately self-consistent under the recurrent dynamics and stable to perturbations orthogonal to its positional mode (see Methods and Appendix).

#### Evidence-driven Edge–Bump Dynamics

We next show that a designed, evidence-dependent modulation of bump–edge feedback (*I*_BE_ and *I*_EB_) enables the network to implement a one-dimensional decision variable governed by a drift–diffusion process.

Assume the feedback input currents between edge and bump are defined by the interaction kernels *W*_BE_ and *W*_EB_, as illustrated in Fig. 2B,

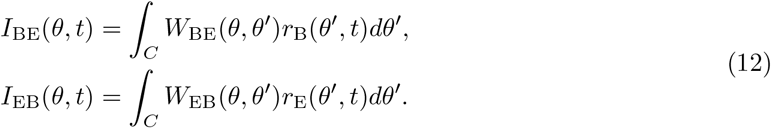

If *I*_EB_ is mediated by a short-range interaction kernel, as is standard in neural-field and continuous-attractor models, the operator *W*_EB_ can be approximated locally by a differential operator acting on the edge activity *r*_E_(*θ*). For a spatially localized and approximately balanced kernel with a dominant antisymmetric component, the leading-order contribution reduces to a first spatial derivative. We therefore choose *W*_EB_ such that

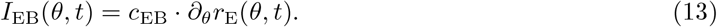

where *c*_EB_ controls the edge-to-bump coupling strength. Since *r*_E_(*θ*) is edge-like, its derivative is a localized bump-shaped profile centered at the edge location. Thus, *I*_EB_(*θ*) provides the bump population with a localized external drive whose center is determined by the edge state (see *I*_EB_(*θ*) in Fig. 2C). A phase offset between the induced input bump and the internal bump state *r*_B_(*θ*) generates tracking dynamics that drives the bump toward the input.

For the bump-to-edge interaction, a centered positive/negative bump input drives the edge right-ward/leftward without deforming its geometry (see *I*_BE_(*θ*) in Fig. 2C). We thereby consider a limiting case of purely local coupling for the bump-to-edge pathway *W*_BE_(*θ, θ*^*′*^) = *c*_BE_*δ*(*θ* − *θ*^*′*^) and thus we have

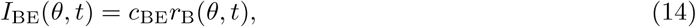

where *c*_BE_ controls the bump amplitude. When the tracking dynamics (modulated by *c*_EB_) is sufficiently fast relative to the intrinsic edge dynamics (modulated by *c*_BE_), the bump effectively remains aligned with the edge location (Fig. 2C). In this aligned regime, the coupled edge–bump state moves approximately as a single mode along the attractor manifold, and its instantaneous velocity is approximately proportional to the bump-to-edge coupling strength *c*_BE_ (see Appendix).

We now parameterize the circuit so that motion along the attractor manifold implements prescribed latent decision dynamics. Because the mapping between the decision variable *x* and the neural position 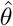 is nonlinear (Fig. 2B, inset), equal displacements in neural space generally correspond to unequal changes in *x*. In the aligned regime, the velocity of the edge–bump state induced by a weak bump-to-edge input is approximately proportional to the local coupling strength *c*_BE_ (see Appendix). By the chain rule, the resulting velocity in decision-variable space therefore scales as

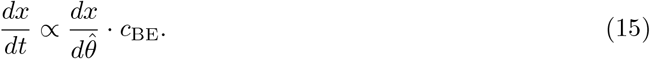

We compensate for this position dependence by letting *c*_BE_ depend on *θ*,

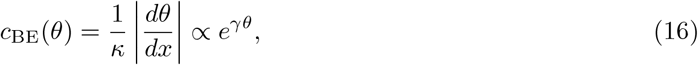

where *κ* is the network-dependent proportionality between bump-to-edge input strength and attractor velocity, and the orientation sign is absorbed into the direction of the scalar drive. With this correction, the same bump-to-edge drive produces an approximately position-independent change in *x*, allowing the circuit input to be matched directly to target behavioral dynamics.

Within the aligned-regime approximation, we next show how the latent decision dynamics can be mapped directly onto the CANN through an appropriate scalar drive. After the geometric correction introduced above, we define the scalar drive *u*(*t*) = *dx/dt*, and modulate the bump-to-edge pathway according to

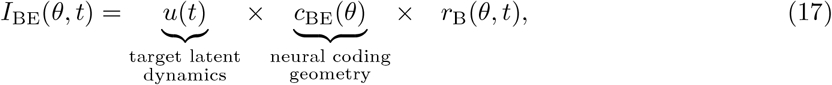

where *u*(*t*) specifies the input required for the decoded CANN state to reproduce the target latent dynamics. Crucially, the coupling profile *c*_BE_(*θ*) depends only on the geometry of the neural representation, whereas the choice of behavioral model enters only through *u*(*t*). Different choices of *f* (*x, t*) can therefore be mapped onto the same CANN without modifying its recurrent architecture.

Notably, this framework does not prescribe a particular neural mechanism by which sensory evidence generates or controls the scalar drive *u*(*t*). For example, in a pulse-based evidence-accumulation task, one could argue that left and right cues selectively gate the bump-to-edge pathways of the corresponding choice-selective groups. Each group would then accumulate evidence favoring its preferred choice, and the latent decision variable would be read out from the difference between the two decoded attractor states. Alternatively, signed evidence could directly determine the direction of the bump-to-edge input, causing both choice-selective groups to encode the same decision variable in complementary coordinates. These implementations are behaviorally equivalent, and *u*(*t*) therefore should be understood as an effective mathematical drive.

### CANN reproduces diffusion decision choice and reaction-time statistics

The construction in the previous subsection provides a circuit-level implementation of any one-dimensional latent decision model by mapping the latent decision variable onto the aligned edge–bump manifold in neural space. To test the framework in a concrete setting, we consider the standard DDM as a special case of the general latent dynamics introduced above

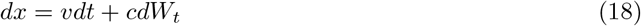

with absorbing boundaries at *x* = 0 and *x* = 1, where *v* is the drift rate, *c* is the noise scale, and *W*_*t*_ is a standard Wiener process. We have the corresponding scalar drive

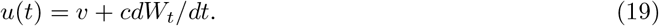

We map this process onto the aligned edge–bump manifold using the geometry-corrected bump-to-edge coupling derived above. This construction predicts agreement at two levels: decoded CANN trajectories should reproduce individual trajectories of the matched DDM, and the resulting choice and reaction-time statistics should agree across trials.

We first show that in a representative condition with *v* = 0.25 and *c* = 0.5, decoded decision-variable trajectories (dashed lines in Fig. 3A, middle) from the circuit closely track the matched diffusion decision trajectories (solid lines in Fig. 3A, middle), and the simulated reaction-time histograms for correct and error trials agree well with the theoretical reaction-time distributions of DDM (Fig. 3A, top and bottom). This agreement extends across a range of parameters. For fixed noise scale *c* = 0.5 and drift rates *v* ∈ [− 1, 1], the quantile-probability functions of the circuit closely overlap those of the matched DDM for both error and correct trials, showing that the model captures the full shape of the reaction-time distributions across conditions (Fig. 3B). The same parameter sweep also yields close agreement in the psychometric curve, with the circuit reproducing the dependence of choice probability on drift rate predicted by DDM (Fig. 3C), and in the mean reaction times, which show the same characteristic dependence on evidence strength as the matched DDM (Fig. 3D).

**Figure 3:**
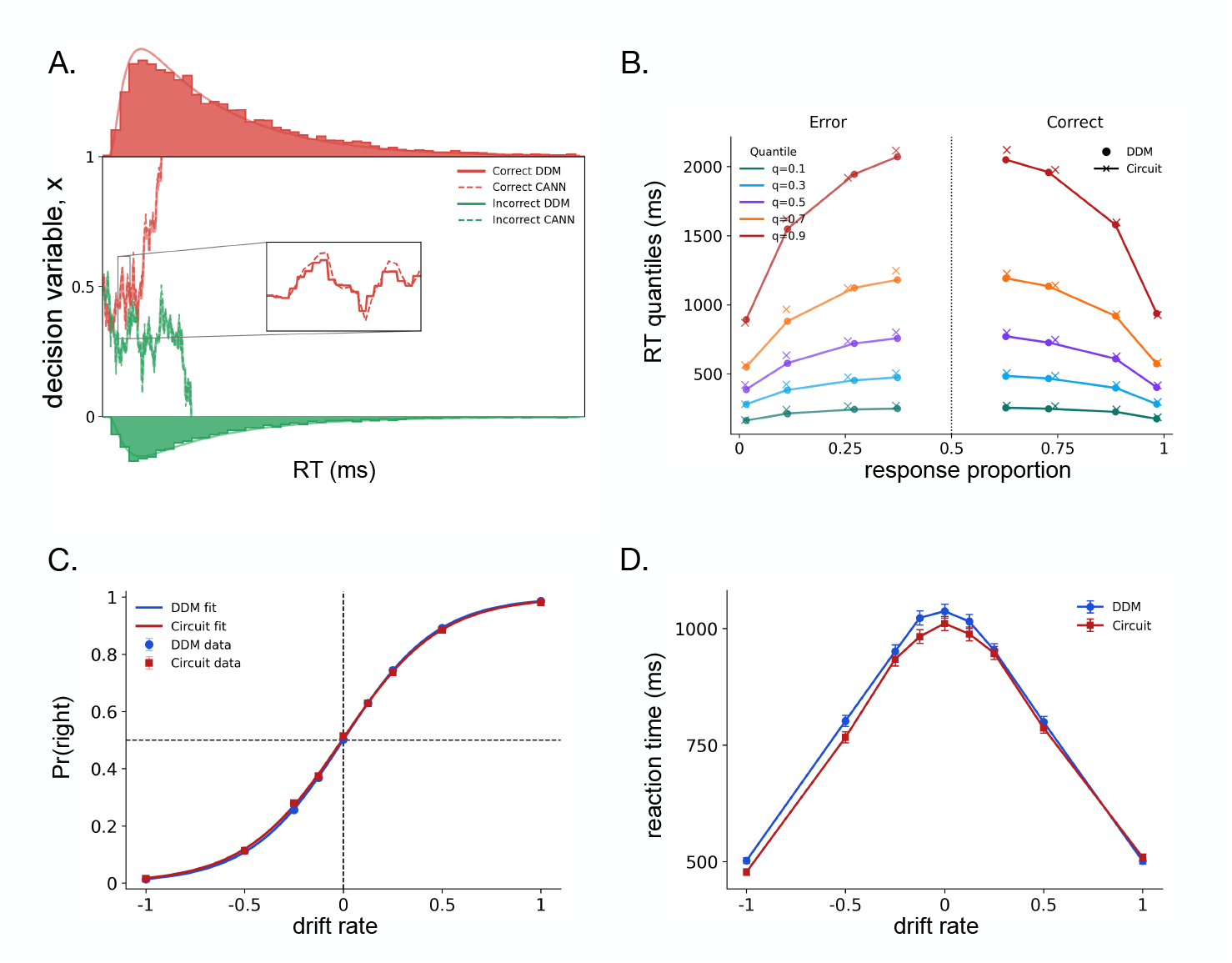
Edge–bump CANN reproduces the behavior of a standard DDM. **(A)** Example decision-variable trajectories across trials as predicted by the matched DDM (solid) and decoded from the circuit model (dashed), for drift rate *v* = 0.25 and noise scale *c* = 0.5. The top histogram represents reaction times for correct trials and the bottom histogram represents reaction times for error trials over 1000 simulations. The solid curves overlaid on the histograms denote the theoretical reaction-time distributions predicted by DDM. **(B)** Quantile-probability functions for error and correct trials under four drift conditions, *v* = ± 0.125, ± 0.25, ± 0.5, and ± 1, with noise scale fixed at *c* = 0.5. Crosses denote circuit-model data and circles denote the corresponding DDM predictions. Across conditions, the circuit model closely tracks the reaction-time statistics of the DDM. **(C)** Psychometric curve showing the probability of a rightward choice as a function of drift rate. The circuit reproduces the dependence of choice probability on evidence strength predicted by DDM. **(D)** Mean reaction time as a function of drift rate. The circuit likewise reproduces the dependence of reaction time on evidence strength across conditions.

Together, these results demonstrate that the designed CANN provides a circuit-level implementation of diffusion decision dynamics. The agreement holds not only for aggregate behavioral measures, such as psychometric curves and mean reaction times, but also for single-trial decoded trajectories and full reaction-time distributions. This establishes an equivalence between the CANN and the matched DDM: the circuit dynamics encodes a one-dimensional variable that obeys the same latent variable dynamics as the DDM.

### CANN generates ramping and sequential neural dynamics

Having matched the latent dynamics and behavioral statistics of a standard DDM, we next examined how the circuit expresses the diffusion decision process at the level of individual neural activity.

We first visualized how a single noisy decision trajectory is represented by the coupled edge–bump circuit. In Fig. 4A, the red trajectory marks the decoded decision variable as it moves through the edge and bump population states. Slices through neural space at fixed times show the instantaneous population profiles: when the decision variable revisits similar values at different times (*t*_1_ to *t*_4_ in fig. 4A), the corresponding edge and bump profiles are nearly identical. This shows that the population states depend on the current decision-variable value rather than elapsed time. Complementary slices at fixed neural positions (*θ*_1_ to *θ*_4_ in fig. 4A) give the activity time courses of individual neurons. These traces show how movement along the shared manifold is expressed locally: edge neurons produce graded ramp-like responses with different ramping speeds, whereas bump neurons fire transiently.

**Figure 4:**
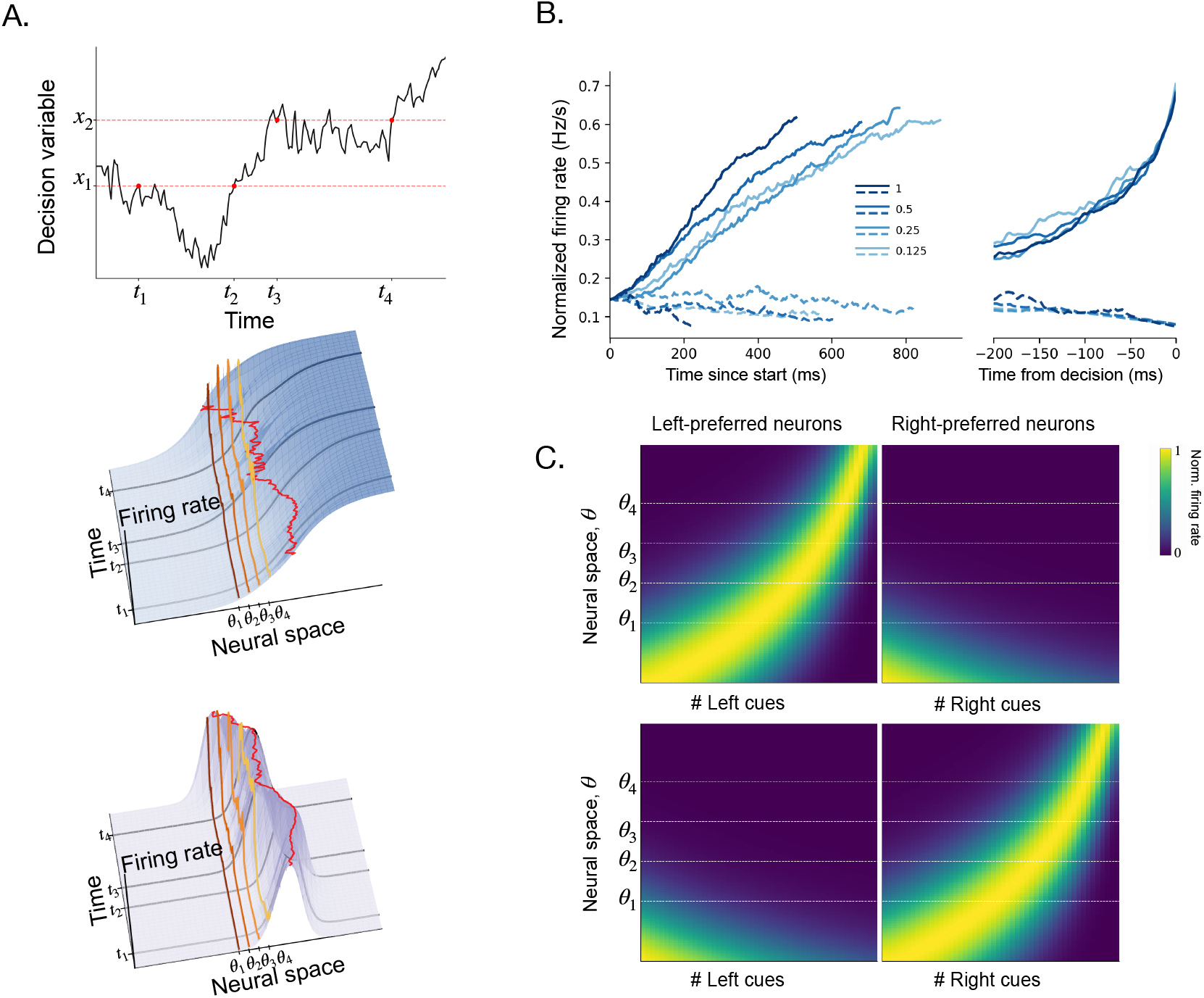
Edge–bump dynamics transform decision-variable trajectories into heterogeneous neural responses. **(A)** Single-trial simulation showing how the circuit represents a noisy decision trajectory. Top, the simulated decision variable evolves over time and revisits similar values at different moments (*t*_1_/*t*_2_ and *t*_3_/*t*_4_). Bottom, the corresponding simulated edge and bump population states represent this trajectory as movement through a shared neural state space. Population profiles at matched decision-variable values are nearly identical despite occurring at different times, showing that activity is decision-variable dependent. Highlighted curves at different neural positions (*θ*_1_ to *θ*_4_) show heterogeneous temporal responses of individual edge and bump neurons, while the red trace marks the encoded decision-variable trajectory through the population. **(B)** Trial-averaged activity of 61 neurons from the edge population across four task difficulties, grouped by drift rate and choice. Each trace is shown until the mean reaction time for its corresponding condition. Positive-drift, bound-reaching trials show ramping activity whose rise is faster for larger drift rates, whereas activity associated with the opposite choice remains low or decreases, illustrating drift-dependent and choice-specific ramping dynamics in the edge population. **(C)** Bump population activity under all-left-cue and all-right-cue conditions, designed analogously to cue-sequence manipulations used to test choice-specific sequences in PPC [14]. Trials were simulated with cues consistently favoring either the left or right bound, and bump neurons were grouped by their preferred choice. The resulting activity shows ordered sequential activation within the bump population, with distinct sequences recruited for left- and right-preferring neurons under the corresponding cue condition. This demonstrates that the bump population expresses the latent decision trajectory as choice-selective sequential activity.

We next asked whether the edge population reproduces the ramping dynamics commonly observed during evidence-accumulation tasks. Empirically, decision-related neurons often show gradual buildup activity that is choice-selective, depends on evidence strength, and reaches commitment earlier on easier trials [4–6,9]. Fig. 4B shows the same qualitative pattern in the model. When trials are grouped by drift rate and choice, activity associated with the selected bound ramps upward, with stronger drift rates producing steeper rises and shorter reaction times. Activity associated with the opposite choice remains weak. Thus, the edge population converts the latent diffusion process into the choice-specific and difficulty-dependent ramping responses observed in neural recordings.

Moreover, cortical recordings during extended evidence-accumulation tasks have revealed neurons with transient, sequential activity, such that different cells are preferentially active at different stages of the evolving decision process [13–15]. To study whether the bump population recovers the same organization, we binned the population bump activities by the decoded decision variable and estimated each bump neuron’s receptive field across this latent coordinate. The resulting receptive fields are localized and tile the decision-variable axis (Fig. 4C), showing that the bump population expresses the latent diffusion trajectory as an ordered sequence of transiently active cells.

## Discussion

We proposed a circuit-level mechanistic account for widely observed heterogeneous neural responses during evidence accumulation, including choice-selective ramping and sequential firing patterns. In our Laplace diffusion decision model, ramping and sequentially firing neurons are not treated as signatures of separate computations. Instead, they form two aligned population codes for the same decision state. Ramping neurons have exponential receptive fields over evidence, with rate constants arranged exponentially across neural position (2), giving rise to an edge-like population profile. Sequentially firing neurons have localized receptive fields over evidence, with centers arranged logarithmically across neural position (9), giving rise to a bump-like population profile. These edge and bump profiles define families of translationally invariant population states, in which accumulated evidence is represented by the aligned edge-bump center. We further constructed a coupled continuous attractor model in which reciprocal interactions move the aligned edge–bump state along this manifold, implementing evidence integration as translation in neural space. Thus, the model connects three levels of description for evidence accumulation: a low-dimensional behavioral model, a population-level coding principle, and a large-scale recurrent architecture.

### Reciprocal coupling as a flexible mechanism for evidence integration

A central challenge for circuit models of evidence accumulation is to explain how heterogeneous neural dynamics convert momentary evidence into a persistent update of the decision state. Continuous attractor models have long provided circuit mechanisms for maintaining and integrating continuous variables, such as head direction [21], eye position [28], or spatial location [24]. In these systems, integration is implemented by inputs that translate the network state along the attractor manifold. Brown et al. [29] recently adapted this logic to evidence accumulation with the position-gated bump attractor model: accumulated evidence is represented by the center of the bump (similar to our bump population), and dedicated shifter neurons receive input from the current bump location and drive the bump to move in response to left or right cues, analogous to rotation cells in head-direction systems. This mechanism requires the cue-activated shifter population to be appropriately registered to the current bump location, so that the bump is pushed to shift towards the input. This mechanism is natural for head-direction systems in the sense that shifter neurons have a clear computational and biological interpretation: they transform angular velocity into a displacement of a circular heading representation. However, the evidence variable in a decision task is often task-defined rather than anatomically fixed. The same circuit may need to accumulate different evidence in different contexts, and a circuit that requires dedicated shifter neurons for each evidence dimension may therefore be less adaptable. To address this challenge, we implemented evidence integration through a reciprocal coupling mechanism between two parallel attractors: the bump provides cue-modulated input to the edge, causing it to move freely along the evidence space, and the bump is continuously registered to the edge by consistently receiving centered-aligned input from the edge. Without requiring an additional cue-specific shifter circuit, this framework therefore allows any form of external input to be reduced to a scalar drive that gates the bump-to-edge pathway and reuses the same coupled attractor architecture to integrate different forms of momentary evidence.

### Logarithmically compressed neural representations

A central assumption of this circuit mechanism is the logarithmic compression of evidence tunings: neural receptive fields for the decision variable are evenly spaced on a logarithmic scale, with preferred evidence values of the n-th neuron satisfying *x*_*n*_ ∝ *e*^*γn*^ (see (1), (2), (8), and (9)). This assumption is motivated by a broader theoretical idea: when a continuous variable spans a wide dynamic range, an efficient population code should allocate receptive fields approximately uniformly in the logarithm of that variable. Such scale-invariant coding has been proposed for time, space, and sound pressure that vary across multiple scales [17, 30–32]. Empirically, logarithmic compression has been widely observed in neural representations of internally generated time, including both temporal context cells with a spectrum of time constants and time cells whose fields tile elapsed time on a compressed scale [33–35]. Recordings during evidence accumulation provide partial support for analogous structure: heatmaps constructed by sequentially firing neurons sorted on their peak time display a characteristic “hook” (as in Fig. 4C) [13, 15] . For ramping neurons, logarithmic compression predicts a power-law distribution of rate constants. Recent recordings have reported non-uniform distribution of rate constants in FOF and ADS [11], but whether their distribution follows this predicted form remains an open question.

### A unified computational scheme for monotonic and non-monotonic tunings

An important implication of the present framework is that monotonic ramping tunings and non-monotonic unimodal tunings need not reflect separate evidence-accumulation mechanisms. Brown et al. proposed two candidate circuits to explain distinct evidence coding schemes across brain regions: competing-chain models produce broad monotonic tuning, whereas position-gated bump attractors produce localized, non-monotonic tuning [29]. Our framework offers a complementary interpretation in which these tuning classes arise from two aligned population codes for the same latent decision variable. The edge population gives rise to monotonic ramping responses, whereas the bump population gives rise to localized sequential responses. Moreover, heterogeneity within each class reflects the internal organization of the corresponding code, with edge neurons carrying different ramping rate constants and bump neurons tiling the evidence axis with different preferred evidence values and field widths.

This unified interpretation is consistent with emerging multi-region views of evidence accumulation. Recent simultaneous recordings across cortical and subcortical structures suggest that distributed neural activity may be better described by a shared accumulator than by independent regional accumulators, while still allowing region-specific contributions to diffusion and commitment [36]. In this sense, regional differences in tuning geometry need not imply separate accumulators: they may reflect different parameter regimes or population-code components of a common decision variable. This makes the edge–bump framework directly testable in large-scale simultaneous recordings. On the one hand, one can test whether ramping and sequentially firing neurons co-vary across regions and predict the same trial-by-trial decision state if they reflect a shared accumulator. On the other hand, it predicts the structured tuning distributions: the rate constants of ramping neurons, the preferred evidence values and the field width of sequentially firing neurons should all exhibit a log-uniform distribution.

### Functional implementations of edge-bump decision circuits

The framework is compatible with a multi-region implementation in which edge-like and bump-like codes are carried by different brain areas and upstream evidence or noise signals modulate their interaction within a shared decision circuit. However, such anatomical separation is not required. If edge and bump codes can instead be implemented as functional subpopulations within a local circuit, the same mechanism can account for mixed monotonic and non-monotonic tunings observed within a single recorded population (for example, see [16]), while remaining testable even in experiments that do not simultaneously sample all regions participating in the decision process.

This view is supported by recent work showing that functional attractor components need not map one-to-one onto anatomically isolated modules. For example, Mei et al. [37] found that the zebrafish head-direction system can contain multiple functional rings within the same anatomical scaffold: one ring represents current head direction, while shifter-like rings provide opposing rotational drive that moves the activity bump clockwise or counterclockwise. Analogously, in the edge–bump decision circuit, edge-like and bump-like populations may be identified functionally by their tuning to the latent decision variable rather than by anatomical separation alone. In this implementation, both populations may rely on related recurrent motifs while differing in boundary conditions and cross-population coupling. These factors determine whether a population expresses an edge-like ramping profile or a localized bump-like profile, and how the two profiles remain aligned during evidence accumulation. Thus, this flexibility broadens the empirical scope of the framework: the model can be tested both in multi-region recordings, where edge-like and bump-like codes may be distributed across areas, and in single-region recordings, where they may appear as intermingled functional subpopulations.

## Methods

### Construction of *k*_**E**_ **and** *k*_**B**_

We aim to construct the recurrent connectivities *W*_EE_ and *W*_BB_, from the base kernels *k*_E_ and *k*_B_, such that (6) and (7) are solutions of

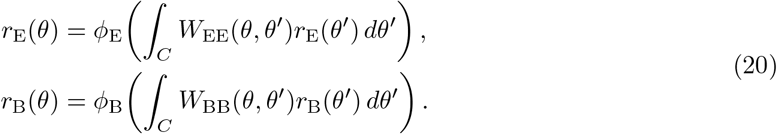

for 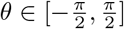.

We start with the edge kernel *k*_E_. We assume that recurrent interactions are translation-invariant in the interior and depend only on neuronal distance, with an even base kernel *k*_E_(Δ). We choose *k*_E_(Δ) from a difference-of-Gaussians (Mexican-hat) family,

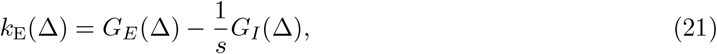

where

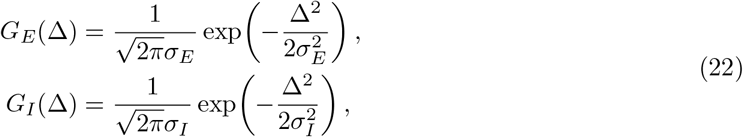

with *σ*_*I*_ = *sσ*_*E*_. We restrict *s* > 1 to enforce local excitation and distal inhibition.

The continuous kernel is discretized over *N* evenly spaced neurons to form a finite recurrent connectivity matrix. Because the simulated neural domain is bounded, we impose a hard-boundary condition on a fraction `clamp_frac` of the neurons and construct the edge connectivity matrix using reflected boundaries: kernel mass extending beyond either end of the finite domain is folded back into the valid domain. After discretization and reflection, we normalize each row of the connectivity matrix to have unit row sum and denote the resulting base matrix by *W*_0_. We then separate the spatial structure of the connectivity from its overall recurrent strength by defining

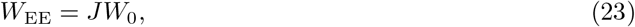

where *J* is a scalar recurrent gain. Reflection and row normalization provide a finite-domain approximation to the translation-invariant recurrent operator while reducing spurious edge drift near the boundaries.

Because the desired edge solutions *ρ*_E_ are sigmoidal, we use the sigmoid nonlinearity

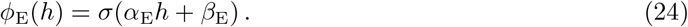

This choice makes approximate self-consistency convenient to enforce in logit space: near the edge transition, the recurrent input should be locally linear in *θ*. Applying the normalized base matrix to the target edge profile gives

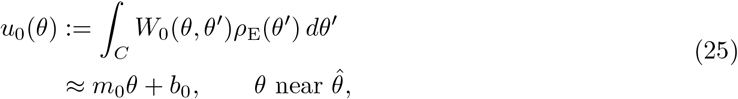

where *m*_0_ and *b*_0_ are obtained from a numerical linear fit in a narrow window around the center of the edge transition.

The target sigmoid edge has slope 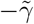 in logit space, whereas the recurrent construction produces a local logit slope *α*_E_*Jm*_0_. We therefore choose

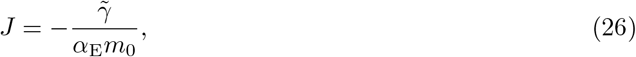

so that the recurrent input matches the desired local slope of the target edge. This calibration ensures approximate self-consistency of the fixed point in the vicinity of the transition, which is the region most sensitive to deviations in position and slope.

Owing to row normalization, the scaled recurrent input

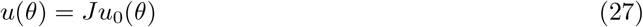

satisfies *u*(*θ*) ≈ *J* in the upper saturated region where *ρ*_E_ ≈ 1, and *u*(*θ*) ≈ 0 in the lower saturated region where *ρ*_E_ ≈ 0. To make the corresponding sigmoid arguments *α*_E_*u* + *β*_E_ approximately symmetric about zero, we set

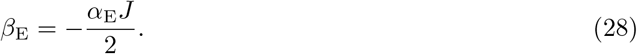

With this choice, the inputs to the nonlinearity are approximately symmetric about zero in the two saturation limits, supporting the global consistency of *ρ*_E_(*θ*) as a fixed point under hard-boundary conditions.

The construction of *k*_B_ follows previous work on a solvable bump-shaped CANN [27]. We used the same parameter `clamp_frac` to set the effective support of the finite bump kernel. Unlike the construction of *k*_E_, we assume *k*_B_ to be pure Gaussian

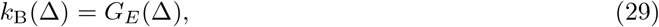

and introduce the square nonlinearity *ϕ*_B_(*h*) = *h*^2^*/C*, where *C* is a constant. Previous studies have shown that CANNs with Gaussian recurrent kernels and quadratic rate nonlinearities are analytically solvable in the continuum limit [27]. Under these conditions, the network admits a continuous family of Gaussian-shaped stationary states with a neutrally stable positional mode (See Appendix for more details).

#### Construction of the position-dependent bump-to-edge coupling

The idealized analysis predicts that the bump-to-edge coupling must vary with position to compensate for the nonlinear relationship between the neural coordinate and the decision variable. In the finite simulation, we decode the edge position using a normalized exponential map over the coding window,

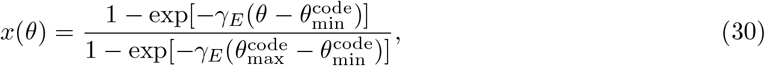

where 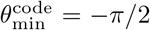 and 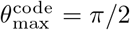. This maps the endpoints of the finite coding window to *x* = 0 and *x* = 1, respectively.

In the weak-input aligned regime, we approximate the edge velocity induced by a constant bump-to-edge input as

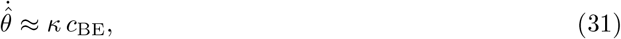

where *κ* is a network-dependent proportionality constant. We estimated *κ* numerically from two constant-coupling calibration runs with *c*_BE_ ∈ {0, 0.04} . For each value, we measured the velocity of the edge position 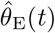 over the valid accumulation window, and estimated *κ* as the slope of the linear relationship between *c*_BE_ and the resulting edge velocity. A single calibrated profile was reused across drift conditions for a given network parameter set.

#### Mapping DDM parameters to circuit drive

After calibrating *c*_BE_(*θ*), the scalar circuit drive was derived directly from the target DDM parameters rather than fitted independently. Under the weak-input aligned approximation, the decoded circuit state obeys

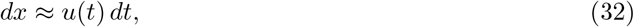

Matching the circuit dynamics to the DDM *dz* = *v dt* + *c dW* gives

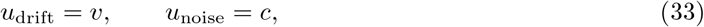

where *v* and *c* are the drift and diffusion parameters of the DDM, respectively. Thus,

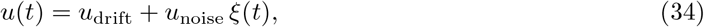

where *ξ*(*t*) denotes standard Gaussian white noise. For the numerical simulations, we discretized this white-noise drive on a time grid with interval Δ*t*_DDM_. For each update window *n*, we sampled an independent Wiener increment Δ*W*_*n*_ ∼ *N* (0, Δ*t*_DDM_) and held the corresponding discretized drive constant within that window:

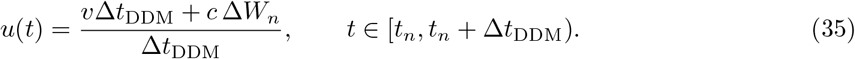

Integrating the held drive over one update window gives Δ*z*_*n*_ = *v*Δ*t*_DDM_ + *c* Δ*W*_*n*_, which matches the increment of the target DDM. Thus, the numerical calibration fits the network-dependent factor *κ* used to construct *c*_BE_(*θ*), whereas *u*_drift_ and *u*_noise_ are set directly by the DDM parameters *v* and *c*, respectively.

#### Network simulation

We simulated the coupled edge–bump network as a finite-dimensional rate model using BrainPy. Each population contained *N* = 1024 units uniformly distributed over *θ* ∈ [− 5*π/*2, 5*π/*2]. The central 20% of this domain, *θ* ∈ [− *π/*2, *π/*2], was used as the coding window for the decision variable, and the outermost 10% of units were clamped to stabilize the finite-domain edge state. The edge recurrent matrix was constructed using the reflected-boundary procedure described above. The bump recurrent matrix was constructed from a row-normalized Gaussian kernel. For the reciprocal pathways, we used a local finite-difference operator for the edge-to-bump input and the identity operator for the bump-to-edge input. Thus, the simulated couplings approximate

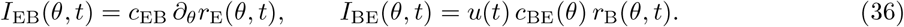

The coding-window position was decoded using Eq. 30. The network was initialized at *x*_0_ = 0.5. We treated *x* = 0 and *x* = 1 as the conceptual absorbing boundaries and terminated a simulated trial when the decoded edge position reached a numerical margin of either boundary (*x* ≤ 0.01 or *x* ≥ 0.99). The scalar circuit drive was updated every 5 ms according to Eq. 34, whereas the intrinsic noise amplitudes of the edge and bump populations were set to zero. The neural rate equations were integrated using BrainPy’s ODE integration routine with a 1 ms simulation step. Evidence input began after a 10 ms initialization period.

For the behavioral comparisons in Fig. 3, we simulated 1000 trials per condition with fixed diffusion scale *c* = 0.5 and drift rates *v* ∈ {0, ± 0.125, ± 0.25, ± 0.5, ± 1}. For the neural-dynamics simulations in Fig. 4, we used the parameter overrides listed in Table 1. The deterministic simulations in Fig. 2 used the same network construction with *u*_noise_ = 0 and a constant positive or negative drift component.

**Table 1:** Baseline network-simulation parameters. These parameters were used unless otherwise stated. For the simulations in Fig. 4, we used *τ*_E_ = 0.3 ms, *c*_EB_ = 0.2, and *σ*_*B*_ = 0.08 to obtain a narrower bump profile and improve tracking of the single-trial neural dynamics.

| Parameter | Value | Role |
| --- | --- | --- |
| $N$ | 1024 | Units per population |
| `coding_frac` | 0.2 | Fraction of neural domain used to encode $x$ |
| `clamp_frac` | 0.1 | Fraction of boundary-clamped units |
| $\tau_{\text{E}}$ | 1 ms | Edge-population time constant |
| $\alpha_{\text{E}}$ | 1 | Edge sigmoid gain |
| $\gamma_{\text{E}}$ | 1.1 | Edge-profile and readout scale |
| $\tau_{\text{B}}$ | 0.15 ms | Bump-population time constant |
| $\beta_{\text{B}}$ | 6 | Bump nonlinearity normalization |
| $\sigma_{\text{B}}$ | 0.2 | Width of the Gaussian bump |
| $c_{\text{EB}}$ | 0.1 | Edge-to-bump coupling strength |
| $\Delta t_{\text{network}}$ | 1 ms | Neural integration step |
| $\Delta t_{\text{DDM}}$ | 5 ms | Drift–diffusion update interval |

## Appendix

### Sigmoid approximation of the edge representation

**Figure S1:**
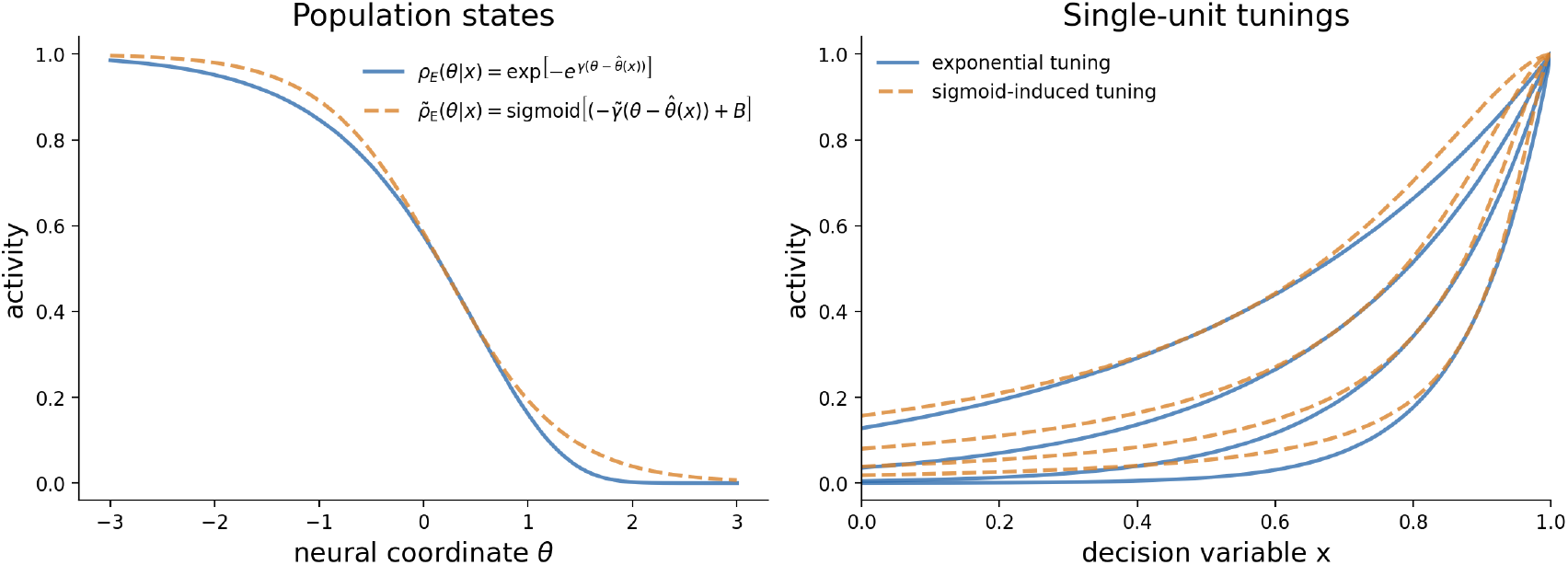
Sigmoid approximation of the edge representation. **Left**, population-level comparison between the double-exponential edge profile induced by exponential ramping tunings (blue) and the biased sigmoid approximation used for analytical tractability (orange dashed). The sigmoid parameters are chosen so that the approximate edge matches both the value and local slope of the original profile at the edge location 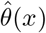. **Right**, single-unit tuning curves obtained by evaluating each population profile at fixed neural coordinates. The sigmoid approximation induces monotonic ramping tunings that closely track the original exponential tunings while preserving the ordering of tuning scales across neurons.

Here we justify the sigmoid approximation used in Eq. (6). The purpose of this approximation is not to match the double-exponential edge profile pointwise, but to preserve the geometric features used by the circuit construction: a monotonic edge, rigid translation with 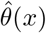, a localized derivative near the edge, and the corresponding ordering of single-neuron ramping tunings.

We choose the bias *B* and slope parameter 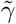 so that the sigmoid edge matches both the value and the local slope of the original edge at 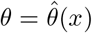. For the double-exponential edge,

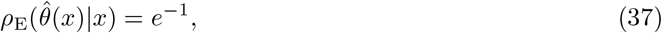

and

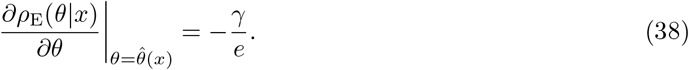

For the sigmoid approximation in Eq. (6), the corresponding value and slope are

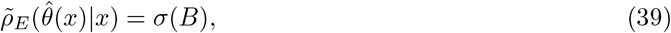

and

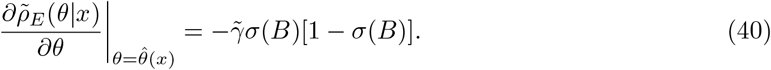

Thus the matching conditions are

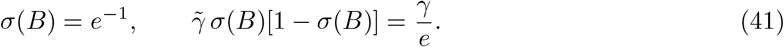

Solving these equations gives

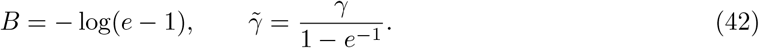

This same approximation also preserves the single-unit ramping structure. For a neuron at position *θ*_*i*_, the exact edge representation induces the single-neuron tuning

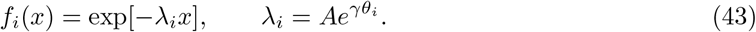

Evaluating the sigmoid approximation at the same fixed neural coordinate gives

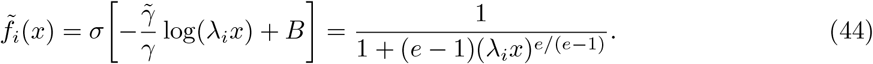

Therefore both the exact and approximate single-unit tunings depend on the same normalized variable *λ*_*i*_*x*. The approximation preserves the monotonic ramping form, the ordering of ramping rates across *θ*_*i*_, and the local value and slope at the characteristic transition point *λ*_*i*_*x* = 1. The remaining discrepancy is a smooth difference in tail shape rather than a change in the represented variable or in the population geometry.

Supplementary Fig. 1 illustrates this approximation at both levels. At the population level, the exact and sigmoid profiles form closely aligned translated edge states across different values of *x*. At the single-unit level, fixed neural positions yield similar monotonic ramping tunings under the exponential and sigmoid-induced representations. Thus, for the stability and positional-dynamics analyses below, the biased sigmoid provides an analytically convenient representative of the same edge code.

#### Stable continuous attractor manifolds for bump and edge populations

Continuous attractor models have classically been used to stabilize localized bump states, such as in heading-direction and path-integration circuits. In edge-bump CANN, we use this standard bump-attractor construction for the sequential population, but we additionally require an edge-like population to behave as a continuous attractor. To support the edge-bump circuit construction, we therefore show below that the two recurrent populations have the required attractor properties: the bump population supports the standard stable family of translated localized states; the edge population supports an analogous family of translated edge states under suitable connectivity.

##### Bump population

We first show that the bump population is a stable solution in the absence of edge-to-bump input.

The bump population obeys

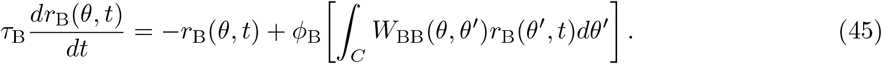

We note that, although the simulated network is defined on a finite neural interval, the bump profile and recurrent kernel are spatially localized, and the coding region is chosen far enough from the boundaries that boundary effects are negligible for the local stability analysis. Since the evidence coordinate is not intrinsically periodic, we therefore approximate the coding domain by the infinite line, *C* ≃ (− ∞, ∞).

A stationary bump solution therefore satisfies

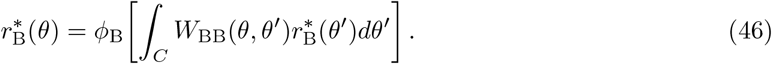

Because *W*_BB_ depends only on distance, any rigid translation of a stationary solution is also stationary. Thus, once one localized bump solution exists, translation invariance immediately generates a continuous family of stationary bump states.

To relate this rate formulation to the standard solvable bump-attractor construction, define the recurrent input

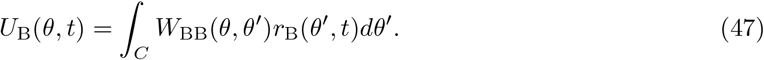

Applying the recurrent operator *W*_BB_ to the rate equation gives

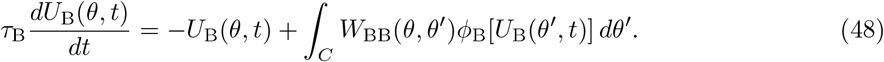

Now our equation has the same form as the dynamical equation in a solvable CANN. If we assume a quadratic nonlinearity 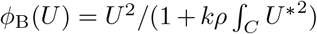, then a stable solution can be analytically written as a Gaussian function

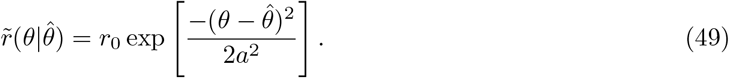

where *r*_0_ depends on the normalized constant *k*, and the bump width *a* depends on the connectivity profile *W*_BB_. We refer the reader to Fung et al. [27] for a detailed proof of stability.

##### Edge population

For the bump population, the Gaussian-kernel and quadratic-nonlinearity construction gives an analytically tractable bump solution. The edge population is less direct: we do not have an analogous closed-form construction that automatically yields the desired sigmoidal edge profile as a stable attractor state. Instead, in the network construction we choose *W*_EE_ so that the target profile *ρ*_*E*_ is approximately self-consistent under the edge dynamics (see Methods).

We therefore analyze here what conditions *W*_EE_ must satisfy for the edge solution 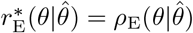 to form a continuous family of stationary states. In what follows, we drop the sub-index E throughout the proof to simplify notation.

Define the translation operator: (*T*_*ϵ*_*f* )(*θ*) = *f* (*θ* − *ϵ*) and assume 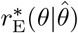 is a fixed point of (10), then any *ϵ*-translation of *r*^*\**^ satisfies

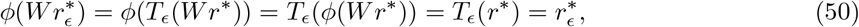

where we use the fact that the translational operator and convolutional operator are commutative. This shows that 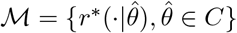 forms a continuous family of fixed points. We thereby choose 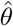 such that 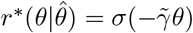 and only consider this solution in what follows. Consider a slight perturbation around the stationary state *r*(*θ, t*) = *r*^*\**^(*θ*) + *δr*(*θ, t*). Using the identity *σ*^*′*^(*h*) = *σ*(*h*)(1 − *σ*(*h*)), we obtain

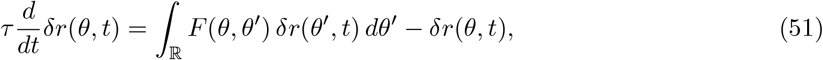

where the interaction kernel is

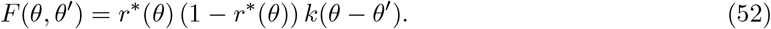

Now we define the linear operator ℒ,

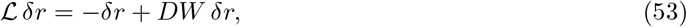

where

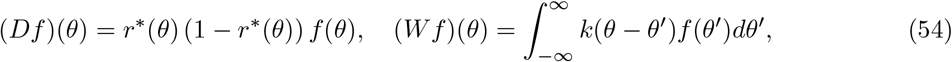

and rewrite the system as

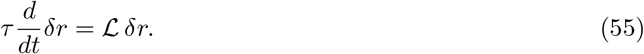

Consider the eigenfunction: then *λ* follows:

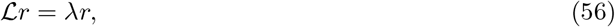

then *λ* follows:

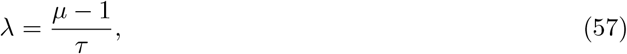

where *µ* is an eigenvalue of *DW* .

The existence of a continuous family of fixed points implies that perturbations tangent to this manifold neither grow nor decay. The corresponding eigenfunction *r*_0_ that satisfies ℒ*r*_0_ = 0 is given by the derivative of the fixed point with respect to its position parameter, evaluated at 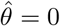:

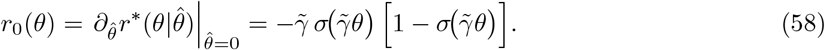

It remains to show that all nontrivial eigenvalues of *DW* lie below 1. Since *k* is even, *W* is self-adjoint on *L*^2^(ℝ). The symmetrized operator

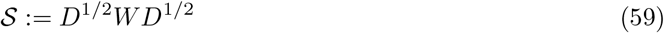

is self-adjoint and similar to *DW* (hence has the same eigenvalues {*µ*_*k*_ }). Because *S* is Hilbert–Schmidt, its eigenvalues satisfy

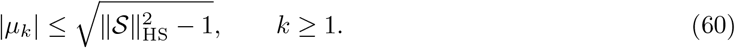

Moreover, by Cauchy–Schwarz,

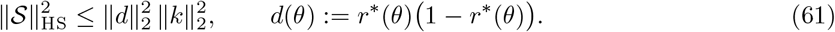

For the sigmoid edge 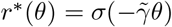 we can compute 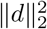 explicitly,

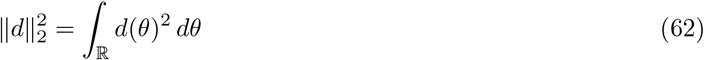

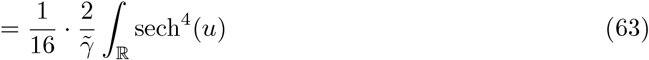

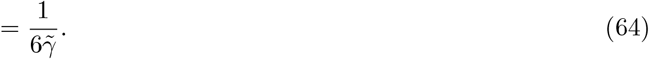

Hence

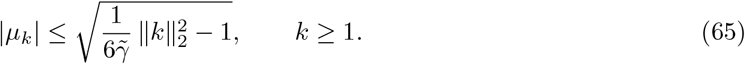

For the difference-of-Gaussians kernel 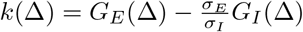,

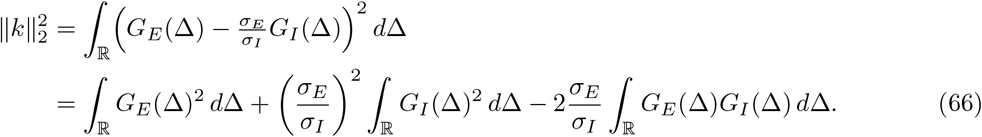

Using the standard Gaussian identities

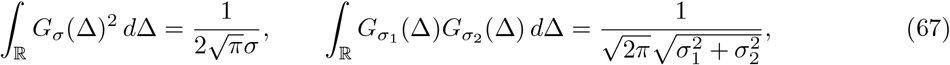

we obtain

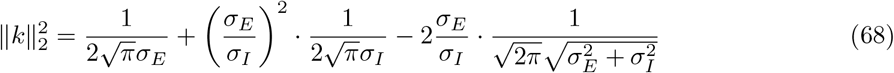

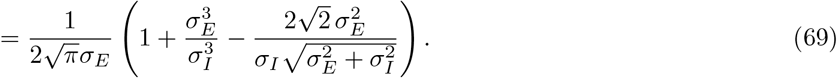

In particular, if the parameters *σ*_*E*_, *σ*_*I*_ are chosen s.t. 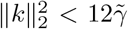, then |*µ*_*k*_| < 1 for all *k* ≥ 1, and therefore the linearized eigenvalues satisfy *λ*_*k*_ = (*µ*_*k*_ − 1)*/τ* < 0.

#### Translations of edge and bump with reciprocal inputs

After showing that the bump and edge populations each support stable translated manifolds, we next examine how reciprocal inputs move the two states along their respective manifolds. We first derive a generic projection formula for the velocity induced by a weak input anchored at a fixed offset from the current attractor state. We then apply the same formula to the two reciprocal pathways: *I*_BE_ sets the edge velocity, whereas *I*_EB_ pulls the bump toward the edge position. The aligned regime corresponds to the case in which the bump can track the edge with a small, approximately constant phase offset.

##### Generic projection onto the positional mode

Consider a population with stable translated states 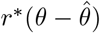 and dynamics

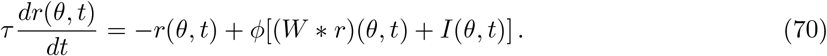

Assume that the population receives a weak input of fixed shape *q* and strength *c* whose center is *θ*_*I*_ (*t*),

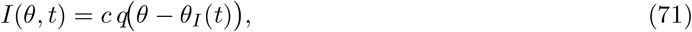

and that the activity remains close to the attractor manifold,

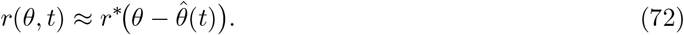

Let 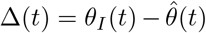 denote the input offset relative to the current state. Linearizing around *r*^*\**^ and projecting the input onto the positional mode gives

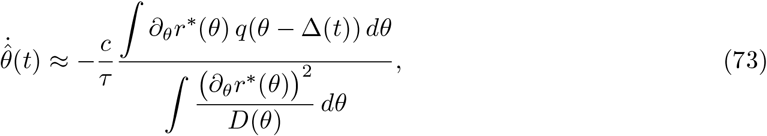

where *D*(*θ*) = *ϕ*^*′*^[(*W* * *r*^*\**^)(*θ*)] is the local gain evaluated at the stationary profile. This expression is the leading-order velocity of the attractor position; higher-order terms correspond to shape deformations, which decay by the stability results above.

##### Edge motion driven by *I*_**BE**_

For the edge population, the bump-to-edge input has the form

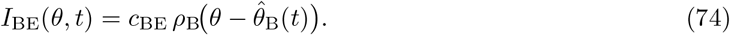

Applying (73) with 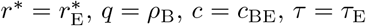, and 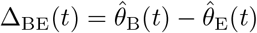 gives

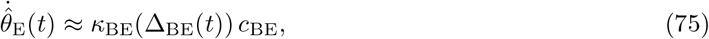

where

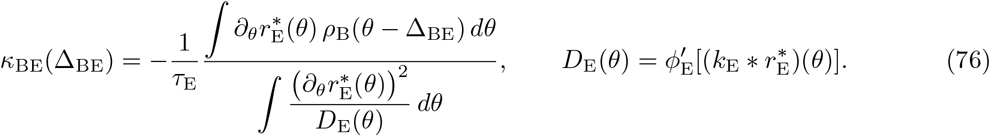

When the bump-to-edge input is approximately centered on the edge transition, Δ_BE_ ≃ 0, this reduces to the speed law used in the main text,

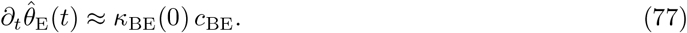

##### Bump tracking driven by *I*_**EB**_

For the bump population, the edge-to-bump input is generated from the spatial derivative of the edge. Up to a sign convention absorbed into *c*_EB_, we write

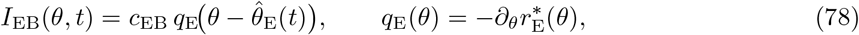

where *q*_E_ is a localized bump-shaped input centered at the edge position. Applying (73) with 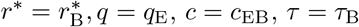, and 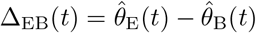gives

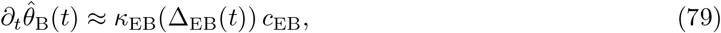

where

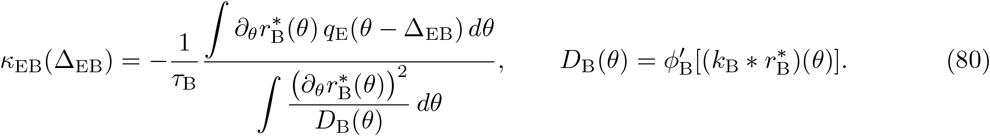

When the edge and bump are exactly aligned, Δ_EB_ = 0, the input is symmetric around the bump center and its overlap with the odd positional mode vanishes. For small offsets, expanding (80) around Δ_EB_ = 0 gives

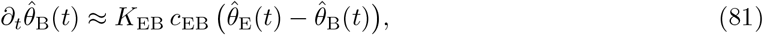

with

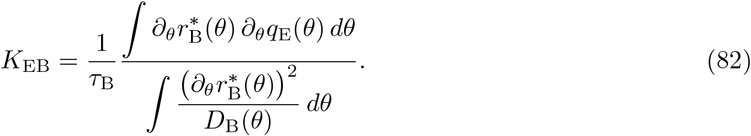

Thus the edge-to-bump pathway acts as a restoring input: the bump remains stationary when it is aligned with the edge, and it moves toward the edge when the edge leads by a small phase offset.

##### Alignment condition

The edge and bump can translate together if there exists a stable phase offset Δ^*\**^ > 0 such that their velocities match. In the constant-coupling reference case, this requires

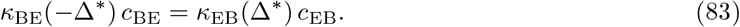

Equivalently, the edge speed induced by *I*_BE_ must be smaller than the maximum tracking speed that *I*_EB_ can induce in the bump,

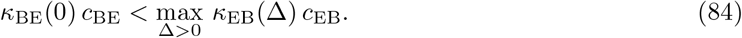

Under the small-offset approximation in (81), the steady offset is approximately

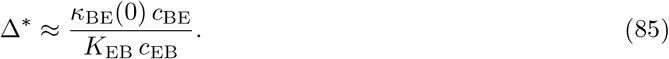

Therefore, when *c*_EB_ is sufficiently large relative to the edge speed set by *c*_BE_, the required phase offset remains small and the coupled edge-bump state moves approximately as a single aligned mode. In the full model, *c*_BE_ varies with position to compensate for the nonlinear decision-variable geometry; the same condition should hold for the largest effective bump-to-edge coupling encountered in the coding region.

## Notes

### Competing Interest Statement

The authors have declared no competing interest.

### Funding

- NIMH: R01MH132171

